# Multidimensional MRI for characterization of subtle axonal injury accelerated using an adaptive nonlocal multispectral filter

**DOI:** 10.1101/2021.07.06.451291

**Authors:** Dan Benjamini, Mustapha Bouhrara, Michal E. Komlosh, Diego Iacono, Daniel P. Perl, David L. Brody, Peter J. Basser

## Abstract

Multidimensional MRI is an emerging approach that simultaneously encodes water relaxation (*T*_1_ and *T*_2_) and mobility (diffusion) and replaces voxel-averaged values with subvoxel distributions of those MR properties. While conventional (i.e., voxel-averaged) MRI methods cannot adequately quantify the microscopic heterogeneity of biological tissue, using subvoxel information allows to selectively map a specific *T*_1_-*T*_2_-diffusion spectral range that corresponds to a group of tissue elements. The major obstacle to the adoption of rich, multidimensional MRI protocols for diagnostic or monitoring purposes is the prolonged scan time. Our main goal in the present study is to evaluate the performance of a nonlocal estimation of multispectral magnitudes (NESMA) filter on reduced datasets to limit the total acquisition time required for reliable multidimensional MRI characterization of the brain. Here we focused and reprocessed results from a recent study that identified potential imaging biomarkers of axonal injury pathology from the joint analysis of multidimensional MRI, in particular voxelwise *T*_1_-*T*_2_ and diffusion-*T*_2_ spectra in human Corpus Callosum, and histopathological data. We tested the performance of NESMA and its effect on the accuracy of the injury biomarker maps, relative to the co-registered histological reference. Noise reduction improved the accuracy of the resulting injury biomarker maps, while permitting data reduction of 35.7% and 59.6% from the full dataset for *T*_1_-*T*_2_ and diffusion-*T*_2_ cases, respectively. As successful clinical proof-of-concept applications of multidimensional MRI are continuously being introduced, reliable and robust noise removal and consequent acquisition acceleration would advance the field towards clinically-feasible diagnostic multidimensional MRI protocols.

## 1 INTRODUCTION

Water molecules within biological tissues interact with their local chemical environment via nuclear relaxation processes and follow diffusion patterns trajectories that are governed by the local tissue density and geometry. Using a combination of magnetic field profiles to probe these mechanisms, magnetic resonance (MR) provides exquisite sensitivity to both the chemical composition, through relaxation parameters, and microstructure, through diffusion parameters, of biological tissues.

One fundamental obstacle for using MRI to characterize tissue heterogeneity is the averaging that occurs across the image volume elements, known as voxels (i.e., pixels with thickness). Voxel-averaged images can only provide macroscopic information with respect to the voxel size, which is typically ~1-3 mm^3^. In a mammalian brain, an individual voxel contains multiple chemical and physical microenvironments such as axons, neurons, glia, myelin, and cerebrospinal fluid. Many biological processes-of-interest take place at a microscopic scale that only affects a small portion of any given voxel, which therefore makes them undetectable using conventional voxel-averaged MRI methods. The inability to separate normal and pathological tissue within a voxel is a major contributor to the insensitivity and ensuing non-specificity of conventional MRI methods in detecting abnormal cellular processes.

By simultaneously encoding multiple MR “dimensions”, such as relaxation times (*T*_1_ and *T*_2_) [1] and diffusion [2, 3], multidimensional distributions of those MR parameters can provide fingerprints of various chemical and physical microenvironments within the volume-of-interest, which can be traced back to specific materials and cellular components. If combined with imaging [4], multidimensional MRI has the potential to overcome the voxel-averaging limitation by accomplishes two fundamental goals: (1) it provides unique intra-voxel distributions instead of an average over the whole voxel; this allows identification of multiple components within a given voxel [5, 6, 7], while (2) the multiplicity of dimensions inherently facilitates their disentanglement; this allows higher accuracy and precision in derived quantitative values [8, 9, 10, 11].

Although traditionally multidimensional MR experiments required many repeated acquisitions and therefore have imposed serious time constraints [12], acquisition strategy [13, 14], computational [15, 6, 3, 16], and pulse design [17, 18] technological breakthroughs have significantly reduced the data burden and positioned multidimensional MRI as a powerful emerging imaging modality for studying biological media. Despite of these advances, wide-spread clinical translation still presents challenges, in particular, due to relatively low signal-to-noise ratio (SNR) and the ensuing increased data amount requirement. To address that, we report the use of a nonlocal estimation of multispectral magnitudes (NESMA) filter [19] on multidimensional MRI data to perform noise reduction for reliable parameter determination and further data reduction. To date, NESMA has been successfully used to improve determination of myelin water fraction from multi-spin-echo MR images [20], or cerebral blood flow from arterial spin labeling MR images [21].

We chose to focus and reprocess a subset of data from our recent study that showed multidimensional MRI can uncover subtle axonal injury patterns in the human brain, otherwise inaccessible using conventional quantitative MRI techniques such as diffusion tensor imaging (DTI), *T*_1_ or *T*_2_ maps [22]. The study investigated brain samples derived from human subjects who had sustained traumatic brain injury (TBI) and control brain donors using MRI, followed by co-registered histopathology that included amyloid precursor protein (APP) immunoreactivity to define axonal injury severity [23]. Abnormal multidimensional *T*_1_-*T*_2_, mean diffusivity-*T*_2_ (MD-*T*_2_), and MD-*T*_1_ spectral signatures that were strongly correlated with injured voxels were identified and used to generate axonal injury biomarker maps [22]. Here we study the effect of applying a multispectral nonlocal filter on three representative cases (a control and two TBI cases), with the main goal of evaluating the performance of NESMA on reduced datasets to limit the total acquisition time required for reliable multidimensional MRI characterization of brain tissue. The co-registered APP histology images serve as a “ground truth” reference, thus providing a unique opportunity to quantitatively evaluate to what extent the accuracy of the injury biomarkers maps is preserved under substantial data reduction.

## 2 METHOD

### 2.1 Donors specimens employed in the present study

We evaluated autopsy-derived brain specimens from two different human brain collections. Formalin-fixed portions of approximately 20×20×10 mm^3^ of the Corpus Callosum (CC) were obtained from one military subject from the DoD/USU Brain Tissue Repository and Neuropathology Program (https://www.researchbraininjury.org, Bethesda, MD; Subject 1), and two civilian subjects enrolled in the Transforming Research and Clinical Knowledge in Traumatic Brain Injury study (TRACK-TBI; https://tracktbi.ucsf.edu/transforming-research-and-clinical-knowledge-tbi) (Subjects 2 and 3). For each case, a next–of-kin or legal representative provided a written consent for donation of the brain for use in research. The brain tissues used have undergone procedures for donation of the tissue, its storage, and use of available clinical information that have been approved by the USU Institutional Review Board (IRB) prior to the initiation of the study. All experiments were performed in accordance with current federal, state, DoD, and NIH guidelines and regulations for postmortem analysis.

Subject 1 was a 44 years old male with no known TBI history and postmortem APP-negative histopathology. Subject 2 was a 60 year old male that died as a result of a intraparenchymal hemorrhage following a motor vehicle accident. Subject 3 was a 49 year old male that died as a result of intraparenchymal and subarachnoid hemorrhages following a fall.

### 2.2 MRI acquisition

Prior to MRI scanning, each formalin-fixed brain specimen was transferred to a phosphate-buffered saline (PBS) filled container for 12 days to ensure that any residual fixative was removed from the tissue. The specimen was then placed in a 25 mm tube, and immersed in perfluoropolyether (Fomblin LC/8, Solvay Solexis, Italy), a proton free fluid void of a proton-MRI signal. Specimens were imaged using a 7 T Bruker vertical bore MRI scanner equipped with a microimaging probe and a 25 mm quadrupole RF coil.

Multidimensional data were acquired using a 3D echo planar imaging (EPI) sequence with a total of 56 and 302 images for *T*_1_-*T*_2_ and MD-*T*_2_, respectively, and with 300 *μ*m isotropic spatial resolution, which resulted in respective acquisition times of 4.5 and 17.8 hr. To test the feasibility of data reduction using NESMA we derived reduced datasets by sub-sampling the full datasets. The total number of *T*_1_-*T*_2_ images was reduced from 56 to 36 (35.7% decrease), while the total number of MD-*T*_2_ images was reduced from 302 to 122 (59.6% decrease). Further details can be found in the Supplementary Material.

The SNR was always maintained above 100 (defined as the ratio between the average unattenuated signal intensity within a tissue region of interest, and the standard deviation of the signal intensity within the background). The sample temperature was set at 16.8°C.

### 2.3 Multidimensional MRI processing

Here we implemented a marginally-constrained, *ℓ*_2_-regularized, nonnegative least square optimization to compute the multidimensional distribution in each voxel, as previously described [8, 24]. It is a well-tested approach that had been proved robust and reliable [25, 26, 27, 2, 28, 29, 14], which in this study had resulted in two types of distributions in each voxel: *T*_1_-*T*_2_ and MD-*T*_2_. The 2D *T*_1_-*T*_2_ and MD-*T*_2_ distributions were evaluated on 50 × 50 logarithmically sampled grids using a previously described algorithm [13]. The range for *T*_1_ was 1 – 10,000 ms, the range for *T*_2_ was 1 – 500 ms, and the range for MD was 0.0001 – 5 *μ* m^2^/ms.

If one considers the multidimensional distributions as spectra, it is possible to use them to generate maps of specific spectral components by means of integration over a predefined parameter range generally associated with a spectral peak. The integral value is a number between 0 and 1, representing a certain spectral component (SC) in a given multidimensional distribution, which can be computed in each voxel to generate an image of that specific SC [30]. Here we apply a recently proposed unsupervised algorithm to identify the injury-associated spectral information [22], and generate injury biomarker maps that closely follow APP histopathology.

### 2.4 The nonlocal estimation of multispectral magnitudes (NESMA) filter

For each sample, the multidimensional distributions were derived from the original multidimensional data as well as from data denoised using the NESMA filter to improve accuracy and precision in derived distributions.

We consider *K* multidimensional images defined on a discrete grid describing the 3D spatial domain spanned by the image. The underlying idea of quantitative filters is to reduce noise by replacing the intensity of a given voxel by an unbiased estimate of its underlying amplitude. This requires selection of voxels that are likely to come from similar tissue. The NESMA filter restores the amplitude, *A*, of an index voxel, *i*, based on intensities of *M* preselected voxels with similar multispectral signal patterns through:

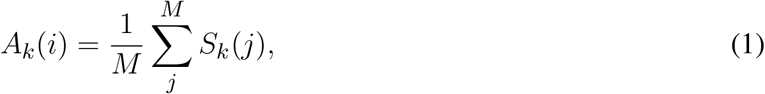

where *S_k_*(*j*) is the measured amplitude in voxel *j* of frame *k. M* is the total number of similar voxels defined using the relative Manhattan distance (RMD) between voxel intensities as

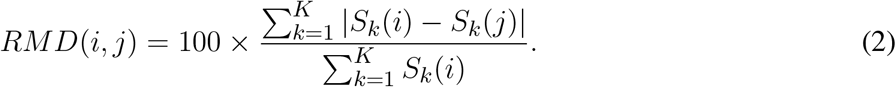

The RMD was calculated between the index voxel *i* and all voxels belonging to a relatively large search window of size *R*, centered around the index voxel *i*, in which emission and reception *B*_1_ fields and noise standard deviation (SD) were assumed to be approximately constant. The size of the window must be sufficiently large to ensure inclusion of an adequate number of similar voxels, and sufficiently restricted to ensure that the transmission and reception *B*_1_ fields and noise SD are approximately constant within the window. In this work, we used a relatively conservative window size to avoid introducing bias in the estimated amplitudes. The size of the search window, *R*, was fixed at 11 × 11 × 11 voxels. Voxels with RMD <5% were considered similar to the index voxel.

### 2.5 Histopathology

After MRI scanning, each CC tissue block was transferred for histopathological processing. Tissue blocks from each brain specimen were processed using an automated tissue processor (ASP 6025, Leica Biosystems, Nussloch, Germany). After tissue processing, each tissue block was embedded in paraffin and cut in a series of 5 *μ*m-thick consecutive sections on which immunohistochemistry for anti-amyloid precursor protein (APP) was performed (DS9800, Leica Biosystems, Buffalo Grove, IL). Further details can be found in [22].

### 2.6 Quantification of axonal damage

Images of APP stained sections were digitized using an Aperio whole slide scanning scanner system (Leica Biosystems, Richmond, IL) at ×20 magnification. The following steps, all implemented using MATLAB (The Mathworks, Natick, MA), were taken to allow for a quantitative analysis of the APP images. First, the images were transformed into a common, normalized space to enable improved quantitative analysis [31]. Then, the normalized images were deconvolved to unmix the primary (APP) and secondary (hematoxylin and eosin, H&E) stains, and background to three separate channels [32]. Once an APP-only image was obtained, a final thresholding step was taken to exclude non-specific staining and to allow for a subsequent % area calculation.

From each tissue section, based on APP staining, traumatic axonal injury (TAI) lesions were identified by an experienced neuropathologist (DI) as white matter (WM) areas with swollen axonal varicosities, axonal bulbs, or distorted axons. Accordingly, regions of interest (ROI) of normal-appearing WM and TAI lesions were manually defined. Additionally, gray matter (GM) ROIs from adjacent cingulate cortex were defined in all sections. Twelve ROIs, covering together an average 81 mm^2^ of tissue, were identified per tissue section. After extracting the ROIs, APP density was expressed as the percentage of total area within the ROI in the binary deconvolved APP image. In total, 36 ROIs from three subjects were included in this study.

## 3 RESULTS

### 3.1 Axonal injury spectral signatures are preserved after filtering

We first investigated the spatially-resolved subvoxel *T*_1_-*T*_2_ and MD-*T*_2_ spectral components to assess the effect of NESMA on the derived voxelwise spectra. To do that, it is useful to summarize the 4D information, which consists of 2D images with 50 × 50 spectra in each voxel, as arrays of images with varying subvoxel *T*_1_, *T*_2_, and MD values. To make them more readable, the 50 × 50 spectra were sub-sampled on a 10 × 10 grid. These maps are shown in Figs. 1, 2, and 3 for all three Subjects. Corresponding histological APP images (co-registered with the MRI) are shown on the left panel of Fig. 4, with red color indicating abnormal APP accumulation.

**Figure 1.**
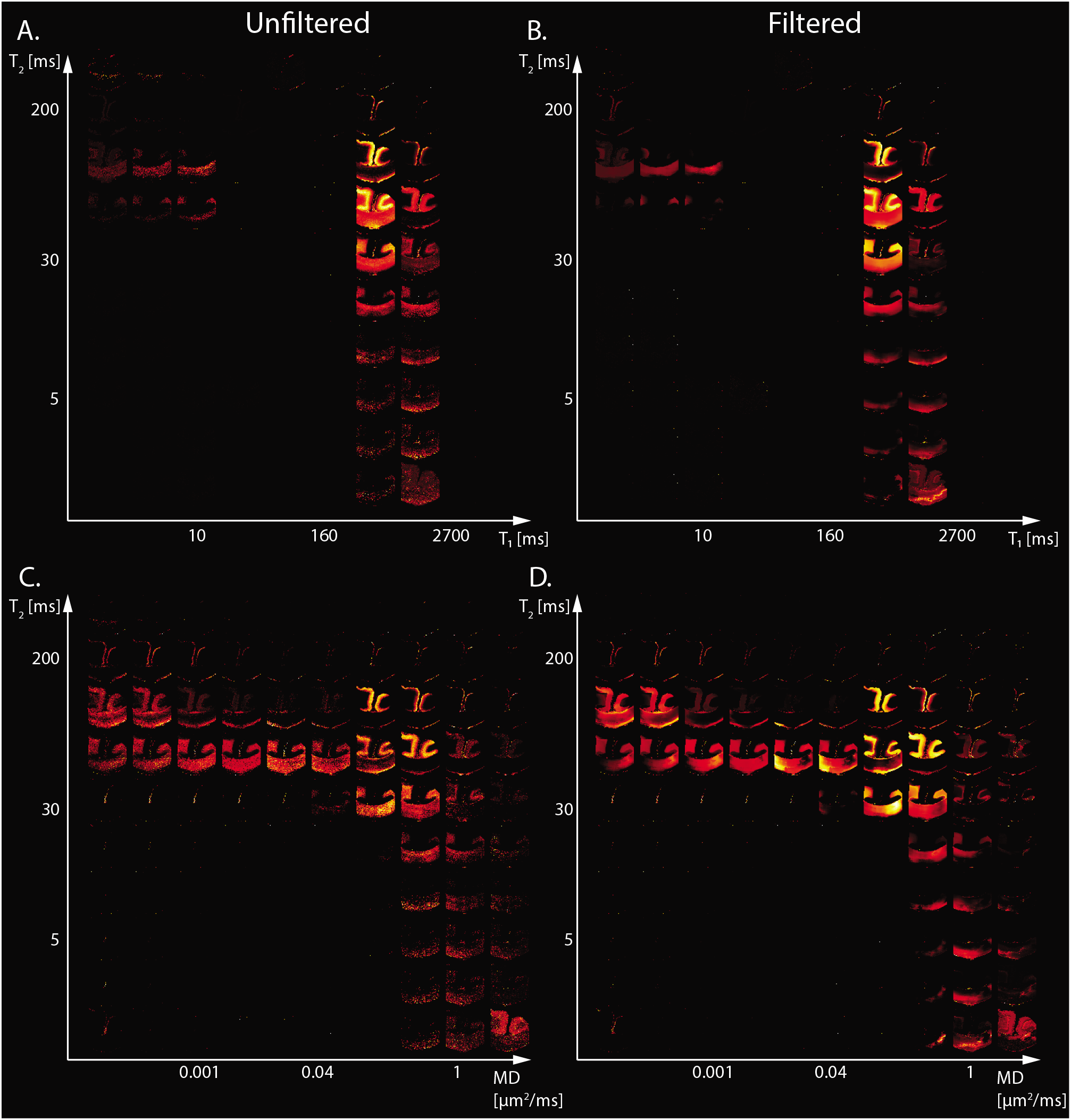
Maps of 2D probability density functions (i.e., 2D normalized spectra) from Subject 1 (control) of (A) unfiltered and (B) filtered subvoxel *T*_1_-*T*_2_ values reconstructed on a 10 × 10 grid of subvoxel *T*_1_ values (horizontal axes) and subvoxel *T*_2_ values (vertical axes), and maps of (C) unfiltered and (D) filtered subvoxel MD-*T*_2_ values reconstructed on a 10 × 10 grid of subvoxel MD values (horizontal axes) and subvoxel *T*_2_ values (vertical axes).

**Figure 2.**
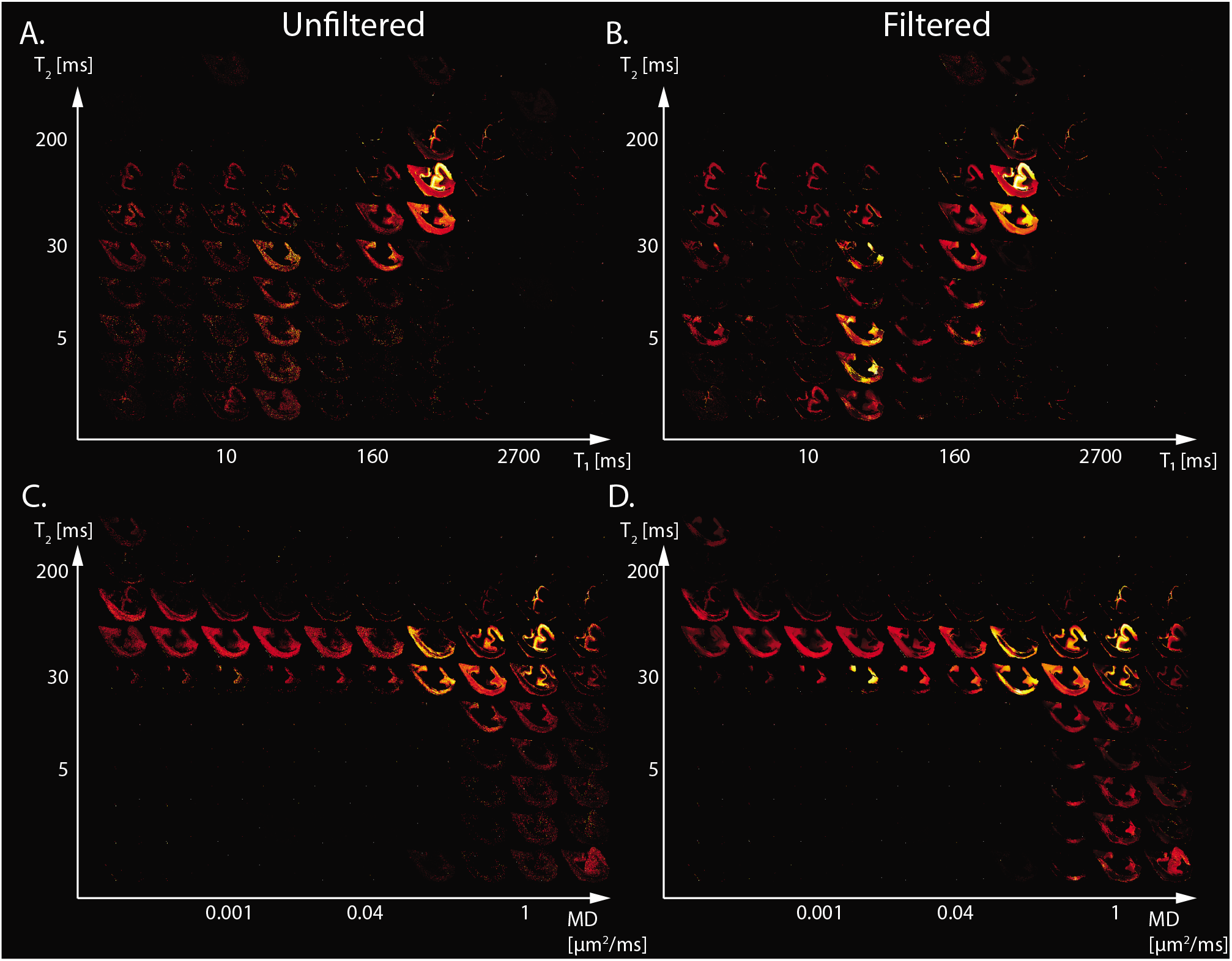
Maps of 2D probability density functions (i.e., 2D normalized spectra) from Subject 2 (TBI) of (A) unfiltered and (B) filtered subvoxel *T*_1_-*T*_2_ values reconstructed on a 10 × 10 grid of subvoxel *T*_1_ values (horizontal axes) and subvoxel *T*_2_ values (vertical axes), and maps of (C) unfiltered and (D) filtered subvoxel MD-*T*_2_ values reconstructed on a 10 × 10 grid of subvoxel MD values (horizontal axes) and subvoxel *T*_2_ values (vertical axes).

**Figure 3.**
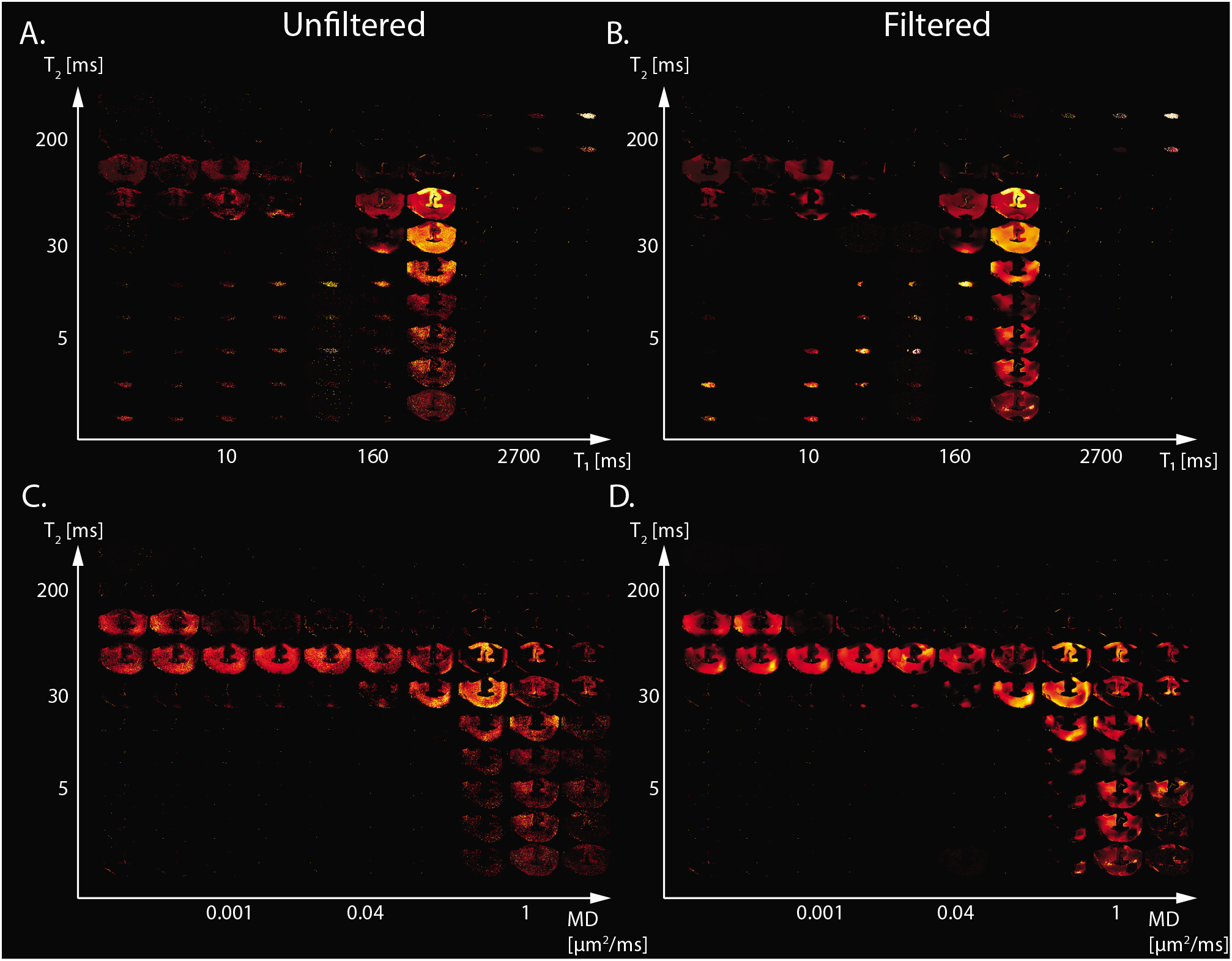
Maps of 2D probability density functions (i.e., 2D normalized spectra) from Subject 3 (TBI) of (A) unfiltered and (B) filtered subvoxel *T*_1_-*T*_2_ values reconstructed on a 10 × 10 grid of subvoxel *T*_1_ values (horizontal axes) and subvoxel *T*_2_ values (vertical axes), and maps of (C) unfiltered and (D) filtered subvoxel MD-*T*_2_ values reconstructed on a 10 × 10 grid of subvoxel MD values (horizontal axes) and subvoxel *T*_2_ values (vertical axes).

**Figure 4.**
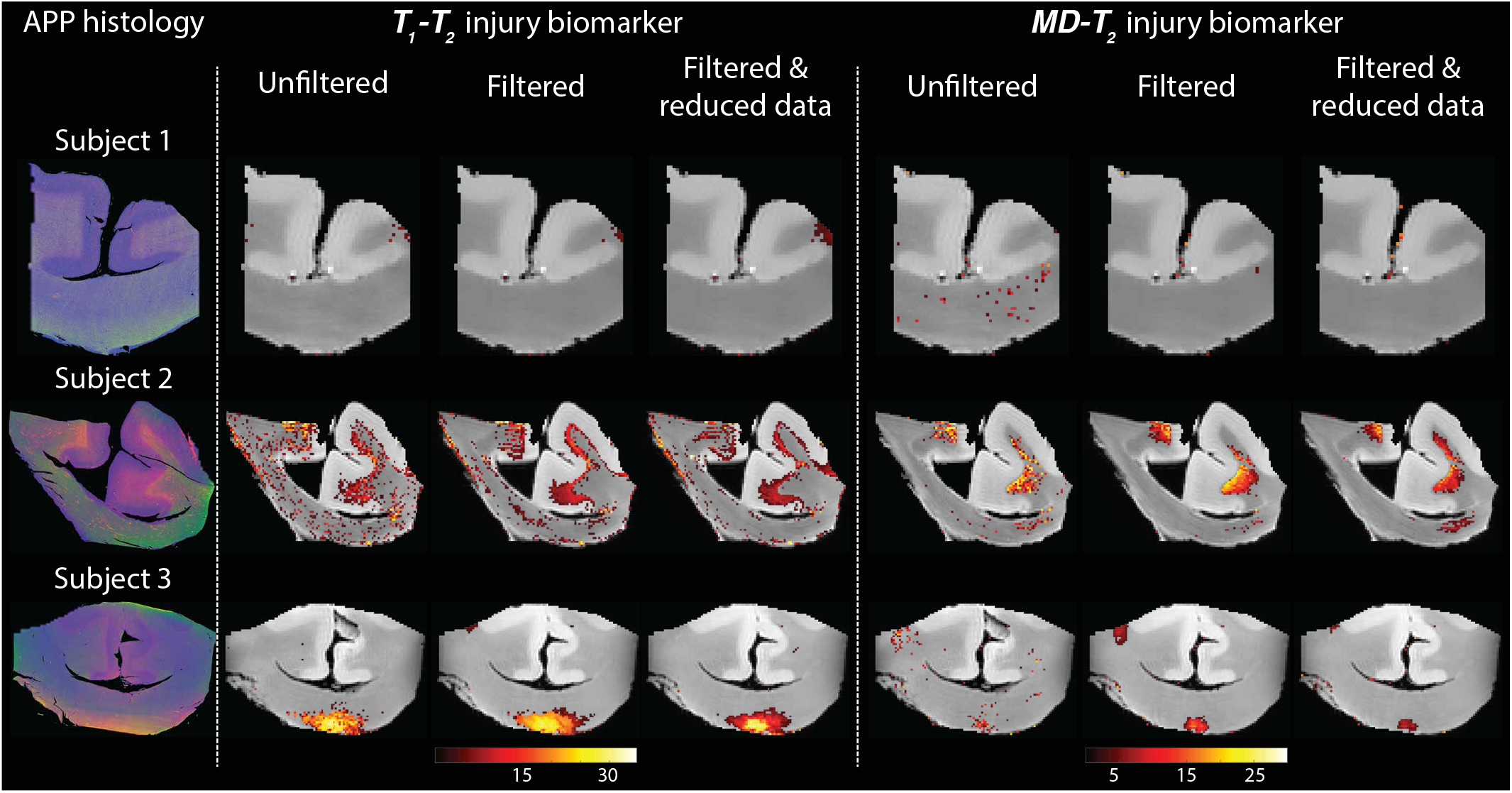
Histological images and multidimensional MR-derived injury biomarker maps of three representative cases, and under different conditions (left to right: unfiltered, filtered, and filtered & reduced data). Deconvolved histological APP images (co-registered with the MRI) are shown on the left panel, red = APP stain (top to bottom: control, and two TBI cases). All multidimensional injury maps were thresholded at 10% of the maximal intensity and overlaid on grayscale proton density images. Multidimensional injury maps of Subject 1 (control) show absent of significant injury under all experimental conditions. Multidimensional injury maps of Subject 2 (TBI) show substantial injury along the white-gray matter interface under all experimental conditions. Multidimensional injury maps of Subject 3 (TBI) show substantial injury at the bottom of the CC under all experimental conditions.

Starting with the control case (Subject 1), the spatially-resolved subvoxel *T*_1_-*T*_2_ and MD-*T*_2_ spectral components are shown in Fig. 1. The left column shows the results from the unfiltered data (*T*_1_-*T*_2_ and MD-*T*_2_ in Figs. 1A and C, respectively), while the right column shows the results from the filtered data (*T*_1_-*T*_2_ and MD-*T*_2_ in Figs. 1B and D, respectively). The maps revealed signal components that were spatially consistent with specific tissue types such as white matter and gray matter.

The spatially-resolved subvoxel *T*_1_-*T*_2_ and MD-*T*_2_ spectral components from the first TBI case (Subject 2) are shown in Fig. 2. As before, the left column shows the results from the unfiltered data (*T*_1_-*T*_2_ and MD-*T*_2_ in Figs. 2A and C, respectively), and the right column shows the results from the filtered data (*T*_1_-*T*_2_ and MD-*T*_2_ in Figs. 2B and D, respectively). Similarly to the control case, here too the maps revealed signal components that were spatially consistent with specific tissue types.

The spatially-resolved subvoxel *T*_1_-*T*_2_ and MD-*T*_2_ spectral components from the second TBI case (Subject 3) are shown in Fig. 3. Unfiltered (*T*_1_-*T*_2_ and MD-*T*_2_ in Figs. 3A and C, respectively) and filtered data (*T*_1_-*T*_2_ and MD-*T*_2_ in Figs. 3B and D, respectively) are shown. As before, signal components that were spatially consistent with specific tissue types as a function of *T*_1_, *T*_2_, and MD were revealed.

Figure 4 shows histological images and multidimensional MR-derived injury biomarker maps of the three representative cases. Histological images (red = APP stain) of the control case (Subject 1) show negative APP staining, compared with positive APP staining in the injured samples (Subjects 2 and 3). We then examine separately the two MRI-derived injury biomarkers, *T*_1_-*T*_2_ and MD-*T*_2_, and show the resulting images obtained using the unfiltered full dataset (as originally published in [22]), the filtered full dataset, and the filtered reduced dataset. In addition, the MRI-derived injury biomarkers obtained by using the unfiltered reduced dataset are shown in Fig. S1 in the Supplementary Material.

Visual inspection of the different injury biomarker maps shown in Figs. 4 and S1 revealed that filtering of the data does not result in loss of the spectral information of interest, and furthermore, the filtered images appear qualitatively of higher quality. Importantly, the data reduction in the case of the filtered data did not significantly affect the resulting injury biomarker maps (Fig. 4).

### 3.2 Evaluation of performance and correlation with histology

Evaluation of filtering performance was based upon the extent of noise reduction and feature preservation, and was quantified by computing the structural similarity index (SSIM) values [33] between the injury biomarker maps under the different experimental conditions (e.g., unfiltered, filtered) and the co-registered APP density histological image as reference. All of the SSIM values are shown in Fig. 5. In the context of the current study we are most interested in the ability to accelerate the multidimensional MRI acquisition, and therefore the accuracy and quality of the reduced data cases are of particular importance. Compared with the unfiltered and reduced data injury biomarker maps, the SSIM values of the filtered and reduced data images increased by 11.1%, 0.9%, and 14.3% for the MD-*T*_2_-based biomarker for Subjects 1 to 3, respectively, and increased by 8.6%, 7.7%, and 4.6% for the *T*_1_-*T*_2_-based biomarker for Subjects 1 to 3, respectively. All of these increases in SSIM were statistically significant (*p* < 0.001).

**Figure 5.**
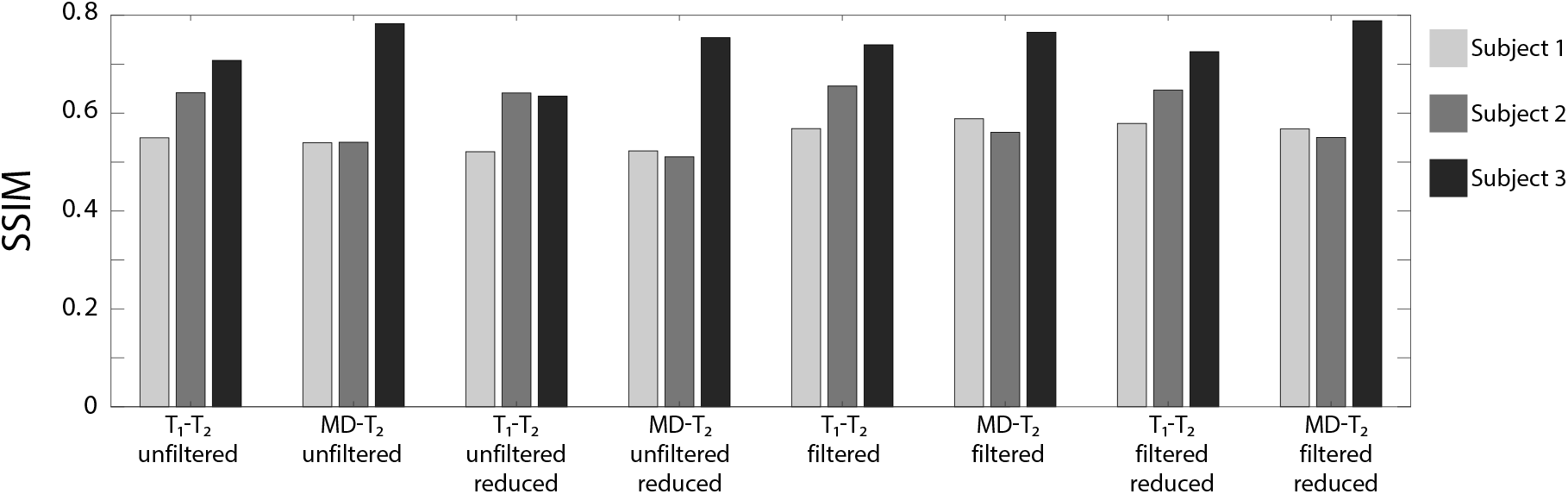
The structural similarity index (SSIM) values between the injury biomarker images under the different experimental conditions (e.g., *T*_1_-*T*_2_ unfiltered, MD-*T*_2_ filtered reduced data) and the co-registered APP density histological image as reference. The three bars at each condition represent the different Subjects (blue = Subject 1, red = Subject 2, and yellow = Subject 3).

To further evaluate the performance of the NESMA filter and the subsequent data reduction, we performed radiological–pathological correlation analyses with histological APP density and all the investigated MRI parameters under the different experimental conditions (e.g., MD-*T*_2_ unfiltered, *T*_1_-*T*_2_ filtered & reduced). Figure 6 summarizes the association between the investigated MR metrics and the pathological findings in normal WM, cortical GM, and TAI ROIs.

**Figure 6.**
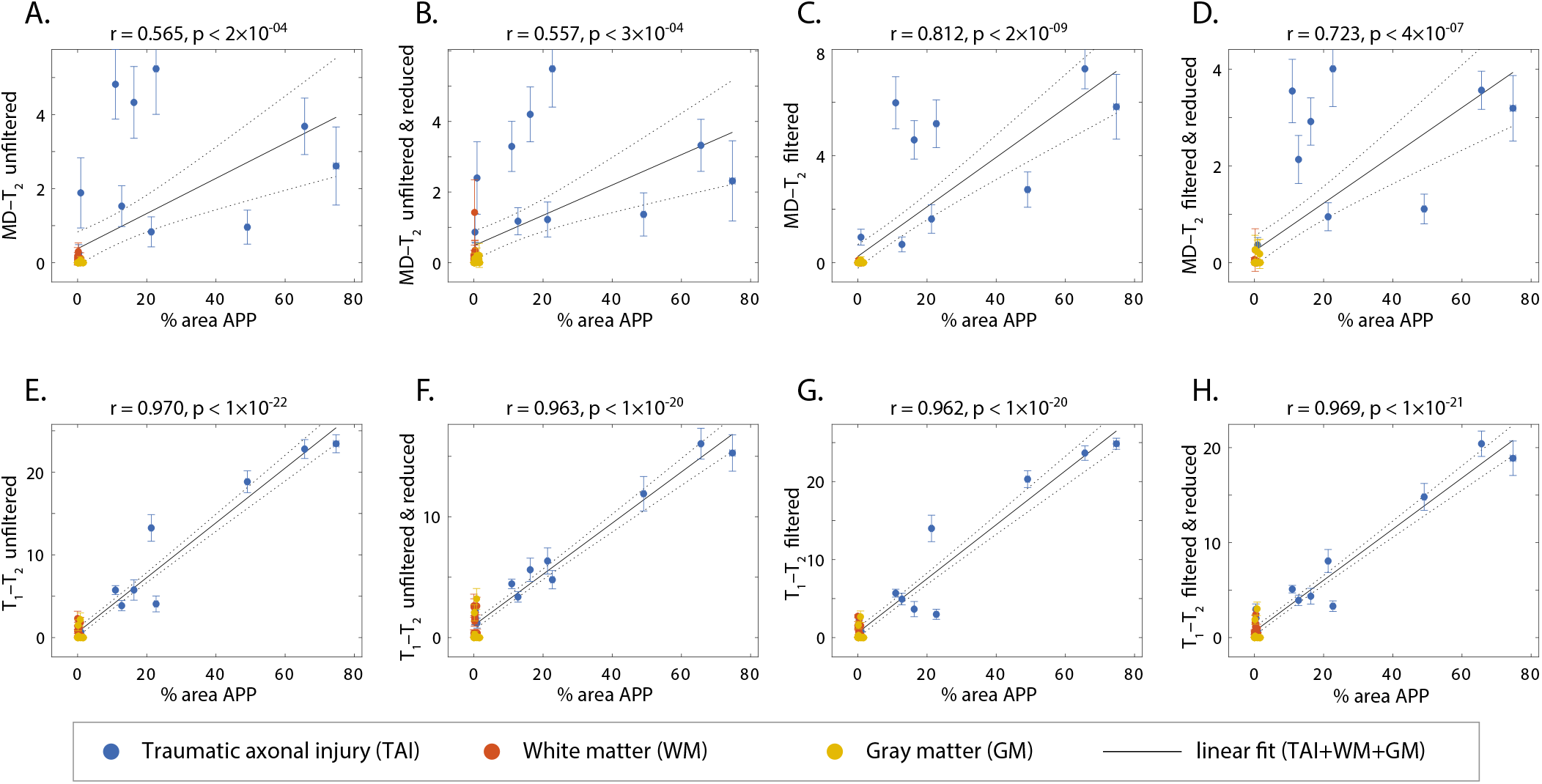
APP density (% area) from 36 tissue regions (APP-positive regions from each TBI case, WM and GM regions), and its correlation with injury biomarker parameter under different experimental conditions. Individual data points represent the mean ROI value from each post-mortem tissue sample. Scatterplots of the mean (with 95% confidence interval error bars) % area APP and (A) MD-*T*_2_ unfiltered, (B) MD-*T*_2_ unfiltered & reduced, (C) MD-*T*_2_ filtered, (D) MD-*T*_2_ filtered & reduced, (E) *T*_1_-*T*_2_ unfiltered, (F) *T*_1_-*T*_2_ unfiltered & reduced, (G) *T*_1_-*T*_2_ filtered, and (H) *T*_1_-*T*_2_ filtered & reduced, show positive and significant correlation with APP density.

To assess the relationship of the MRI parameters with the degree of injury, all tissue ROIs were grouped together and correlated with the APP density (solid lines in Fig. 6). All multidimensional injury biomarker cases under every experimental conditions were strongly and significantly positively correlated with the APP density (i.e., % area APP). These correlations illustrate how the multidimensional injury biomarker maps provide “true negative”, in the sense that any region outside of the TAI lesions has zero or close to zero intensity.

Of particular interest was the case of the MD-*T*_2_ injury biomarker, where the unfiltered dataset resulted in relatively scattered correlation (*r* = 0.565, *p* < 2 × 10^−4^, Fig. 6A), which was largely unaffected by the data reduction (*r* = 0.557, *p* < 3 × 10^−4^, Fig. 6B). Marked improvement was observed after filtering the data, which led to significantly tighter correlation between the MRI biomarker and the histological marker (*r* = 0.812, *p* < 2 × 10^−9^, Fig. 6C). As can be expected, the 59.6% reduction in the data amount led to some reduction in the radiological–pathological correlation (*r* = 0.723, *p* < 4 × 10^−7^, Fig. 6D), although still performing better than the unfiltered full MD-*T*_2_ dataset. Furthermore, filtering led to a reduced variance within the ROIs, as seen by the decrease in the 95% confidence intervals (error bars in Fig. 6).

The *T*_1_-*T*_2_ injury biomarker derived from the unfiltered dataset had excellent correlation with histological APP density (*r* = 0.970, *p* < 1 × 10^−22^, Fig. 6E), and therefore, it is not surprising that this strong relationship was maintained under all of the investigated experimental conditions (*r* = 0.963, *p* < 1 × 10^−20^, *r* = 0.962, *p* < 1 × 10^−20^, and *r* = 0.969, *p* < 1 × 10^−21^, Figs. 6F-H, for unfiltered & reduced, filtered, and filtered & reduced, respectively).

## 4 DISCUSSION

Here we report the use of the NESMA filter on multidimensional MRI data, in particular voxelwise *T*_1_-*T*_2_ and MD-*T*_2_ spectra in fixed human Corpus Callosum, to remove noise and reduce total scan time. We focused on results from a recent study that identified potential imaging biomarkers of axonal injury pathology from the joint analysis of multidimensional MRI and histopathological data [22]. These axonal injury maps were shown to be significantly and strongly correlated with histological evidence of axonal injury. Reprocessing these data provided an opportunity to test the performance of the NESMA filter and its effect on the accuracy of the injury biomarker maps, relative to the histological reference.

Our findings showed that noise reduction in the multidimensional MRI data using an adaptive nonlocal multispectral filter (i.e., NESMA [20]) improved the accuracy of the resulting injury biomarker maps, and furthermore, allowed for data reduction of 35.7% and 59.6% from the full dataset, which led to using only 36 and 122 images in the *T*_1_-*T*_2_ and MD-*T*_2_ cases, respectively.

Specifically, visual inspection and a side-by-side comparison of the unfiltered and filtered subvoxel *T*_1_-*T*_2_ and MD-*T*_2_ spectral components (Figs. 1, 2, and 3) showed that the filtered maps exhibit lower random variations, in particular at the lower ends of the spectra, and that there was no apparent loss of spectral information. For example, Subject 3 exhibited a relatively focal axonal injury at the bottom of the CC (Fig. 4, left panel), captured at the lower end of the *T*_1_-*T*_2_ spectra, which was previously associated with axonal injury [22]. Noticeable noise reduction at these spectral lower ends was observed, which is crucial to the robust identification of axonal injury from these multidimensional MRI data.

Visual inspection of the resulting *T*_1_-*T*_2_ and MD-*T*_2_ injury biomarker maps with respect to co-registered APP histological images suggested improved accuracy after applying the NESMA filter, even after the data was reduced (Figs. 4). Quantitative evaluation that compared the SSIM between the co-registered APP histological images and injury biomarker maps derived from unfiltered and filtered reduced datasets showed a significant increase as a result of the filtering across all subjects (Fig. 5).

We performed radiological–pathological correlation analyses with histological APP density and all the investigated MRI parameters under the different experimental conditions to assess quantitatively whether and to what extent the proposed approach preserves strong correlations even under substantial data reduction (Fig. 6). This analysis indicated that not only the correlations were preserved, but furthermore, they were considerably improved, even after data reduction, as a result of filtering the data. Lastly, our results suggest that the previously proposed [22] adaptive method of locating the injury-associated *T*_1_-*T*_2_-MD spectral signature is robust to noise removal procedures and to data reduction.

Common to all *ex vivo* human MRI studies, our data include the effects of post-mortem degeneration, fixation and resulting dehydration. Because *T*_1_, *T*_2_, and diffusion dynamics are different in fixed tissue compared with living systems, further investigation will be needed to establish whether and how the axonal injury-related *T*_1_-*T*_2_-MD range of multidimensional magnetic resonance parameters is altered in vivo. In this context it is important to note that our findings are not based on absolute values of *T*_1_, *T*_2_, and MD, which indeed are expected to change in vivo. Instead, all of the injury biomarkers maps are generated using the relative signal fraction of an automatically identified *T*_1_-*T*_2_-MD range that does not depend on the actual values of these parameters [22]. We therefore anticipate that *in vivo* spectra will be shifted in all *T*_1_, *T*_2_, and MD dimensions compared to our *ex vivo* findings, however, the distributions of the signal fractions should largely remain similar.

The performance of the NESMA filter does not depend on the particular multidimensional spectra quantification processing pipeline because the filter is applied in the image domain, before its transformed into voxelwise spectra. Here we applied a constrained *ℓ*_2_ regularized inversion framework [13], however, we anticipate that the demonstrated improvement after filtering could be extended to other approaches such as *ℓ*_1_ regularization [34], Monte-Carlo inversion [35, 36], and InSpect [16].

Multidimensional MRI is an emerging approach [37] that is now being applied to address a range of medical conditions such as prediction of pregnancy complications via placenta characterization [9], spinal cord injury [6, 38], prostate cancer [39], breast cancer [40], and axonal injury due to TBI [22]. Recent *in vivo* proof-of-concept applications of subvoxel *T*_1_-*T*_2_ correlation spectra using 105 images [41] and of subvoxel diffusion-*T*_1_ correlation spectra using 363 [11] and 304 [42] images are promising. Here we showed that accurate and robust subvoxel *T*_1_-*T*_2_ and MD-*T*_2_ correlation spectra can be obtained using only 36 and 122 images, respectively, by using a constrained optimization data processing framework (i.e., MADCO [13]) in conjunction with applying the NESMA filter to reduce noise. A reliable and robust noise removal and consequent acquisition acceleration should further advance the field towards clinically-feasible diagnostic multidimensional MRI protocols.

## CONFLICT OF INTEREST STATEMENT

The authors declare that the research was conducted in the absence of any commercial or financial relationships that could be construed as a potential conflict of interest.

## AUTHOR CONTRIBUTIONS

DB: conceptualization, design of the study, methodology, software, investigation, data curation, writing—original draft, writing—review and editing, visualization, supervision, and project administration. MB: conceptualization, design of the study, methodology, software, and writing—review and editing. MK: methodology, investigation, and writing—review and editing. DI: methodology, investigation, and writing—review and editing. DP: investigation, methodology, resources, and writing—review and editing. DLB: design of the study, investigation, resources, and writing—review and editing. PB: conceptualization, methodology, resources, writing—review and editing, and funding acquisition. All authors contributed to the article and approved the submitted version.

## FUNDING

This research was partially supported by a grant from the U.S. Department of Defense, Program Project 308430 Uniformed Services University of the Health Sciences (USUHS). Support for this work also included funding from the U.S. Department of Defense to the Brain Tissue Repository and Neuropathology Program, Center for Neuroscience and Regenerative Medicine (CNRM). DB and MEK were supported by the CNRM Neuroradiology-Neuropathology Correlations Core. MB was supported by the Intramural Research Program of the National Institute on Aging. DI, DPP, and DLB were supported by the CNRM and USUHS. PJB was supported by the Intramural Research Program of the Eunice Kennedy Shriver National Institute of Child Health and Human Development.

## ACKNOWLEDGMENTS

We thank the subjects’ families that consented for brain donations for the better understanding of TBI consequences. The authors thank Mrs Patricia Lee, Mrs Nichelle Gray and Mr Paul Gegbeh for their valuable technical work. We are grateful to Mrs Stacey Gentile, Mrs Deona Cooper and Mr Harold Kramer Anderson for their administrative assistance. We thank the TRACK-TBI Investigators (https://tracktbi.ucsf.edu/transforming-research-and-clinical-knowledge-tbi).

The opinions expressed herein are those of the authors and are not necessarily representative of those of the Uniformed Services University of the Health Sciences (USUHS), the Department of Defense (DOD), the NIH or any other US government agency.

## SUPPLEMENTAL DATA

Supplementary material is available online.

## DATA AVAILABILITY STATEMENT

The datasets generated and analyzed during the current study are available from the corresponding author on reasonable request.

## Notes

### Competing Interest Statement

The authors have declared no competing interest.

### Summary of Updates

Data added to allow a rigorous quantitative analyses (both structural similarity index -SSIM, and radiological-pathological correlations) to support the claims of this study, as well as adding additional details regarding the technical aspects of the NESMA filter.

