## Supplementary Material for "Multidimensional MRI for characterization of subtle axonal injury accelerated using an adaptive nonlocal multispectral filter"

### 1 MRI ACQUISITION

Multidimensional data were acquired using a 3D inversion recovery diffusion-weighted sequence with a repetition time of 1000 ms, and an isotropic voxel dimension of 300 $\mu$m. To encode the multidimensional MR space spanned by $T_1$ and $T_2$ (i.e., $T_1$-$T_2$) and by $T_2$ and diffusion (i.e., MD-$T_2$), the 1D distributions of $T_1$ and $T_2$ were estimated using 20 logarithmically sampled $\tau_1$ (inversion time) values ranging from 12 to 980 ms and 20 logarithmically sampled $\tau_2$ (echo time) values ranging from 10.5 to 125 ms, respectively. For the 1D distribution of MD we used the isotropic generalized diffusion tensor MRI (IGDTI) acquisition protocol to achieve an efficient orientationally averaged diffusion-weighted signal (Avram et al., 2018) with the following parameters: 16 linearly sampled $b$-values ranging from 2,540 to 14,700 s/mm$^2$ in 3 directions, 14 linearly sampled b-values ranging from 4,140 to 14,700 s/mm$^2$ in 4 directions, and 9 linearly sampled b-values ranging from 8,260 to 14,700 s/mm$^2$ in 6 directions, using the efficient gradient sampling schemes in Table 2 in (Avram et al., 2018). This type of diffusion encoding increases the contrast given by local anisotropy and is not intended to measure the isotropic diffusion in the system. Additional diffusion parameters were gradient duration of $\delta = 4$ ms and diffusion time of $\Delta = 15$ ms.

The 2D distributions of MD-$T_2$ and $T_1$-$T_2$, were estimated, respectively, with the following data acquisition protocols (in conjunction with the *a priori* obtained 1D distributions as constraints): A 2D D-$T_2$-weighted data set with 16 sampled combinations of echo times and b-values within the aforementioned 1D acquisition range; and a 2D $T_1$-$T_2$-weighted data set with 16 sampled combinations of inversion and echo times within the aforementioned 1D acquisition range.

To test the feasibility of data reduction using NESMA we derived reduced datasets by sub-sampling the full datasets in the following manner: (1) reducing the 1D encoding data by a factor of two, and in the MD-$T_2$ dataset (2) reducing the number of echo times by a factor of two, and the number of b-values from four to three. The total number of $T_1$-$T_2$ images was therefore reduced from 56 to 36 (35.7% decrease), while the total number of MD-$T_2$ images was reduced from 302 to 122 (59.6% decrease).

### 2 SUPPLEMENTARY FIGURES

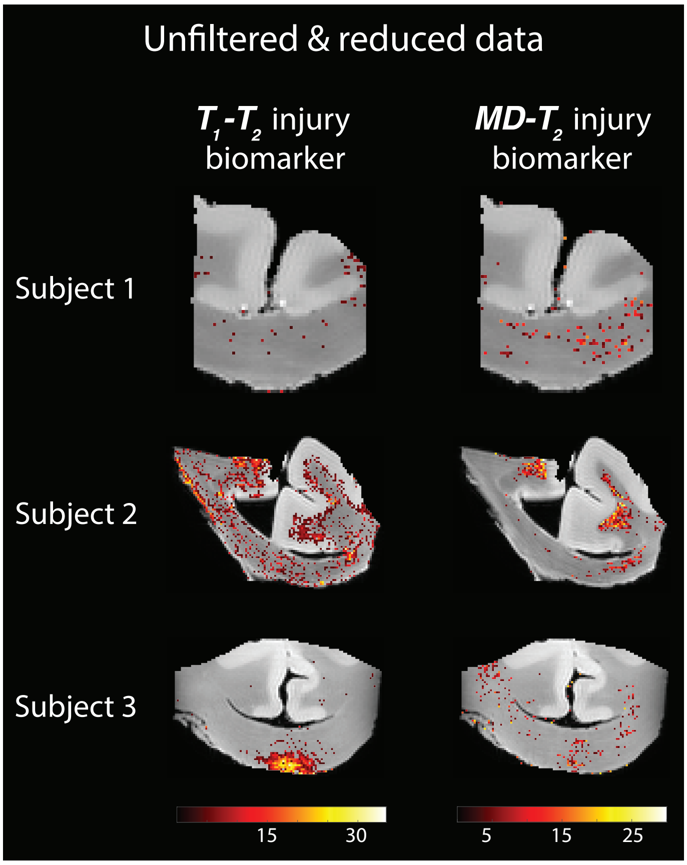


**Figure S1.** Multidimensional MR-derived injury biomarker images obtained using an unfiltered and reduced dataset. All multidimensional injury maps were thresholded at 10% of the maximal intensity and overlaid on grayscale proton density images. Multidimensional injury maps of Subject 1 (control), in particular the MD-$T_2$ biomarker, show spurious signal intensities. Multidimensional injury maps of Subject 2 (TBI) show substantial injury along the white-gray matter interface. The $T_1$-$T_2$ (but not the MD-$T_2$) injury biomarker map of Subject 3 (TBI) shows substantial injury at the bottom of the CC.
